# Native PGC-LC-MS Profiling Reveals Distinct O-Acetylation Patterns of Sialylated N-Glycans Across Mammalian Sera

**DOI:** 10.1101/2025.10.19.683285

**Authors:** Christopher Ashwood

## Abstract

*O*-acetylation of sialic acids represents an additional layer of structural diversity and biological complexity, occurring at various hydroxyl positions (commonly C-7, C-8, or C-9) of the sialic acid residue. This modification modulates the recognition of sialylated glycans by lectins, antibodies, and viral proteins, and contributes to viral tropism and host susceptibility, particularly in influenza and coronaviruses that bind *O*-acetylated sialylated receptors. However, current LC-MS glycomics workflows commonly employ reduction or permethylation, which, while improving chromatographic stability and ionisation, result in the loss of labile *O*-acetyl groups, obscuring their biological relevance. Native glycan analysis, in contrast, preserves the complete structural integrity of glycans, enabling accurate detection of labile modifications.

Using a native released glycan workflow limited to pH ≤ 8, *O*-acetylated *N*-glycans were detected in mouse and rat sera that were previously undetectable under basic derivatisation conditions. Beam-type collision-induced dissociation generated the most informative fragmentation spectra, with diagnostic ions confirming *O*-acetylated NeuGc and NeuAc residues. Chromatographic profiling revealed later elution and broadened peak shapes for *O*-acetylated species, consistent with increased hydrophobicity and microheterogeneity. A checkpoint-based identification workflow incorporating isotopic, chromatographic, and MS2 criteria reduced false positives, retaining only 3-5% of putative *O*-acetylated glycans as confident identifications. Quantitative comparison across species revealed extensive *O*-acetylation in rat (53.4%) and moderate modification in mouse (8.8%), but none detectable in human serum. These findings establish a robust analytical framework for native detection and characterisation of *O*-acetylated *N*-glycans, revealing species-specific regulation of this labile modification.

## Introduction

Sialylation is a critical modification commonly found at the terminal positions of glycans and glycolipids, where it plays a key role in modulating cellular interactions^1^. The sialic acid family consists of nine-carbon α-keto sugars that can occur in multiple structural forms and through diverse glycosidic linkages, contributing to the overall complexity of sialylation. Positioned at the outermost ends of glycan branches, sialic acids influence molecular recognition, cell signaling, and protein stability^2^. Aberrations in sialylation have been associated with numerous pathological states, including cancer, inflammation, and neurodegenerative disorders. Beyond these disease contexts, sialic acids play essential roles in biological processes such as immune regulation^3^, cell signaling^4^, and pathogen recognition^5^.

*O*-acetylation of sialic acids represents an additional layer of structural diversity and biological complexity. This modification occurs at various hydroxyl positions (commonly at C-4, C-7, C-8, or C-9) on the sialic acid residue^6^. *O*-acetylation of sialic acids can modulate the recognition of sialylated glycans by lectins, antibodies, and viral proteins. Viruses, in particular influenza and coronaviruses, are capable of utilizing the *O*-acetylated sialylated glycans found in the respiratory epithelium as receptors^7^. Determining factors in host susceptibility and viral tropism requires investigation into O-acetylation patterns.

The labile nature of sialic acids and their modified *O*-acetylated counterparts pose unique analytical challenges. Current liquid chromatography-mass spectrometry (LC-MS) methods often employ chemical derivatisation steps such as reduction^8^ or permethylation^9^ to chromatographically stabilize the glycan and improve ionization efficiency of glycans, respectively. While these methods are robust for general glycan analysis, they can cause loss of *O*-acetyl groups, thereby precluding the detection of *O*-acetylation altogether^10^. This challenge is compounded by the fact that glycans which lose labile *O*-acetylation become indistinguishable from those that were never *O*-acetylated, making it difficult to confirm the true absence of this modification.

To preserve and accurately characterize labile modifications, native glycan analysis (pH < 8) is essential for preserving the *O*-acetyl modification. This mild pH retains the structural integrity of the glycan, enabling the detection of naturally occurring modifications and allowing for biologically relevant interpretations. With these glycans intact, they are then suitable for characterisation by analytical methods for structural elucidation including LC, ion mobility, and fragmentation. MS-based fragmentation in particular is a promising approach, as it enables structural interpretation without requiring standards^11^.

Upon collision induced dissociation (CID), glycans produce characteristic fragment ions corresponding to specific monosaccharide losses, linkage cleavages, and cross-ring fragments. The interpretation of these fragment masses enables the identification of branching patterns, linkage positions, and monosaccharide composition, thereby allowing detailed structural elucidation^12^. This fragment-based approach circumvents the need for authentic standards, which are often unavailable for complex or rare glycans, and instead relies on the predictable and interpretable nature of glycosidic and cross-ring fragmentation patterns to achieve confident structural assignment. While quadrupole isolation of specific precursor m/z values is possible, a form of separation prior to MS detection is needed to ensure isomer discrimination such as porous graphitised carbon (PGC) LC.

PGC-LC is a powerful technique for the separation of glycans based on their isomeric and linkage-specific structural features. Typically, glycan reduction is employed prior to PGC analysis to remove these anomeric artifacts, which are irrelevant to biological glycan structures but can complicate chromatographic interpretation^8^. However, as shown by Ashwood, *et al*^13^, optimising column temperature, pH, and mobile phase composition, eliminates the anomeric effect on PGC without the need for chemical reduction. This advancement enables native PGC-LC-MS analysis, allowing for the direct, isomer-specific separation of *O*-acetylated *N*-glycans while preserving their native modifications and avoiding split anomeric peaks.

Here, we apply a native PGC-LC-MS workflow to *N*-glycans released from mammalian sera. By limiting sample preparation to pH ≤ 8, *O*-acetylated *N*-glycans were detected across mouse and rat *N*-glycans and fragmentation energy optimised to develop a checkpoint-based identification workflow incorporating isotopic, chromatographic, and MS2 criteria to reduce false positive identifications of *O*-acetylation. Finally, we apply confident *O*-acetylated NeuAc and NeuGc identifications to human sera, ruling out detectable *N*-glycan *O*-acetylation.

## Methods

### Serum and chemical sources

All chemicals and reagents were purchased from Millipore Sigma (Castle Hill, Australia) unless specified otherwise. Sera (human, rat, and mouse) were purchased from Millipore Sigma (Castle Hill, Australia).

### Enzymatic N-glycan release

Sera were solubilised in 5% SDS and 100 mM TEAB. Proteins were reduced with 5 mM dithiothreitol for 30 min at 55 °C, alkylated with 10 mM iodoacetamide in the dark for 30 min at room temperature, and quenched with an additional 5 mM dithiothreitol for 15 min at room temperature. Glycans were released from glycoproteins and purified for LC-MS analysis as described in Ashwood *et al*^*14*^. In brief, proteins were precipitated with methanol and phosphoric acid, and purified via DNA miniprep silica columns (Bioneer, Republic of Korea). Immobilised protein on the miniprep silica columns was resuspended in 160 µL of 50 mM TEAB (pH 8) and 50 units of PNGase-F (Promega, Wisconsin, USA) and left to react at 37 °C overnight.

Following glycan release, glycans were eluted from the silica columns with 400 μL of 0.1% formic acid and then desalted using Supelclean ENVI-Carb SPE (100 mg). The desalting procedure involved conditioning with 400 μL acetonitrile/0.1% formic acid, equilibration with 1.5 mL water/0.1% formic acid, sample loading, desalting with 1.5 mL water/0.1% formic acid, and final elution with 400 μL 50:50 acetonitrile:water containing 0.1% formic acid. The desalted glycan solutions were then dried by centrifugal evaporation. Glycans were resuspended in ultra-pure water and then transferred into a 96 well PCR plate for injection. For reduced N-glycan analysis, glycans were dried by centrifugal evaporation after elution from silica columns, then reduced by 1 M NaBH_4_ in 50 mM KOH and incubated for 3 hours at 50 °C.

### LC-MS setup

Glycans were separated with a Thermo Fisher Scientific Vanquish Horizon HPLC (San Jose, USA) and ionised into an Orbitrap IQ-X Tribrid mass spectrometer (San Jose, USA). A Thermo Fisher Hypercarb PGC column (Lithuania, 100 mm length by 1 mm internal diameter, 3 micron pore size), held at 90 °C, was used for all separations. Mobile phase A composed of water and mobile phase B composed of acetone with 5 mM HFIP and 5 mM butylamine added. LC separation was performed at 150 μL/min. Glycans were separated over a 30 min run, with 0-25% B over 23 min, 100% B held for 3 min, then 100% A for 4 min.

The IQ-X was set to negative mode in DDA MS2 mode with a total cycle time of 1.3 seconds. ESI voltage was 2.8 kV, with sheath and auxiliary gases at 30 and 20 arbitrary units, respectively. Precursor spectra were collected by a full scan of 337 – 1700 *m/z* in an Orbitrap at 60,000 resolution to hit an AGC target of 1e^6^ with a maximum inject time of 400 ms.

Resonant CID and beam-type fragment spectra were collected with an isolation width of 1.5 m/z, maximum injection time of 200 ms, and an AGC target of 1e^5^. The normalised collision energy (NCE) was set to 37 and scanned at 0.6 m/z resolution in a linear ion trap. Dynamic exclusion was utilised, excluding each precursor for 6 seconds after fragmentation. For collision energy optimisation experiments, a targeted precursor list was utilised and MS2 were collected at 25, 32, 37, 42, and 47 NCE with a maximum injection time of 75 ms.

### Data analysis

LC-MS raw files were analysed using GlyCombo^15^ (v1.2) to assign glycan compositions to precursor m/z values, identify the most intense MS2 scans for each structure for annotation in GlycoWorkBench^16^, and construct the Skyline assay detailing *N-*glycan compositions. mzML input was used with an error tolerance of 25 ppm, reducing end specified as free, derivatisation as native and adducts set as M-H^-^. Monosaccharide search space was: Hex 4-12, HexNAc 2-8, dHex 0-1, NeuAc 0-3, NeuGc 0-3, and three custom monosaccharides (NeuGc-Ac) and (NeuAc-Ac) at 0-2 of each. The monoisotopic mass and chemical formula were assigned for each modification as follows: NeuGc-Ac (C_13_H_19_N_1_O_10_, 349.10090 Da), and NeuAc-Ac (C_13_H_19_N_1_O_9_, 333.10598 Da). Additional acetylation was added post-search in Skyline and subsequently quality filtered.

Skyline-daily^17,18^ (v25.1) was used to integrate the first three isotopic peaks with mass analyser set to centroid at 15 ppm mass accuracy. These isotopic integrations were used to quality filter identifications (>=0.85 idotp) and quantify glycans. The idotp value of 0.85 was empirically selected to remove poor quality MS1-matches (caused by monoisotopic peak misassignment, incorrect charge assignment, and poor signal to noise ratios) while preserving high-quality matches^19^.

## Results and discussion

PGC-LC-MS analysis of reduced mouse serum *N*-glycans revealed profiles broadly consistent with previously reported reduced mouse *N*-glycans, including the predominant bi-antennary structure bearing two NeuGc residues. However, under native conditions with a maximum pH of 8 (specifically during PNGase-F release), several low-intensity *O*-acetylated NeuGc-containing glycans were detected that were previously undetectable under the more basic conditions used to generate reduced alditol-bearing *N*-glycans (**Figure 1**). The presence of these minor acetylated forms highlights that conventional reduction and permethylation workflows can obscure labile modifications such as *O*-acetylation.

**Figure 1.**
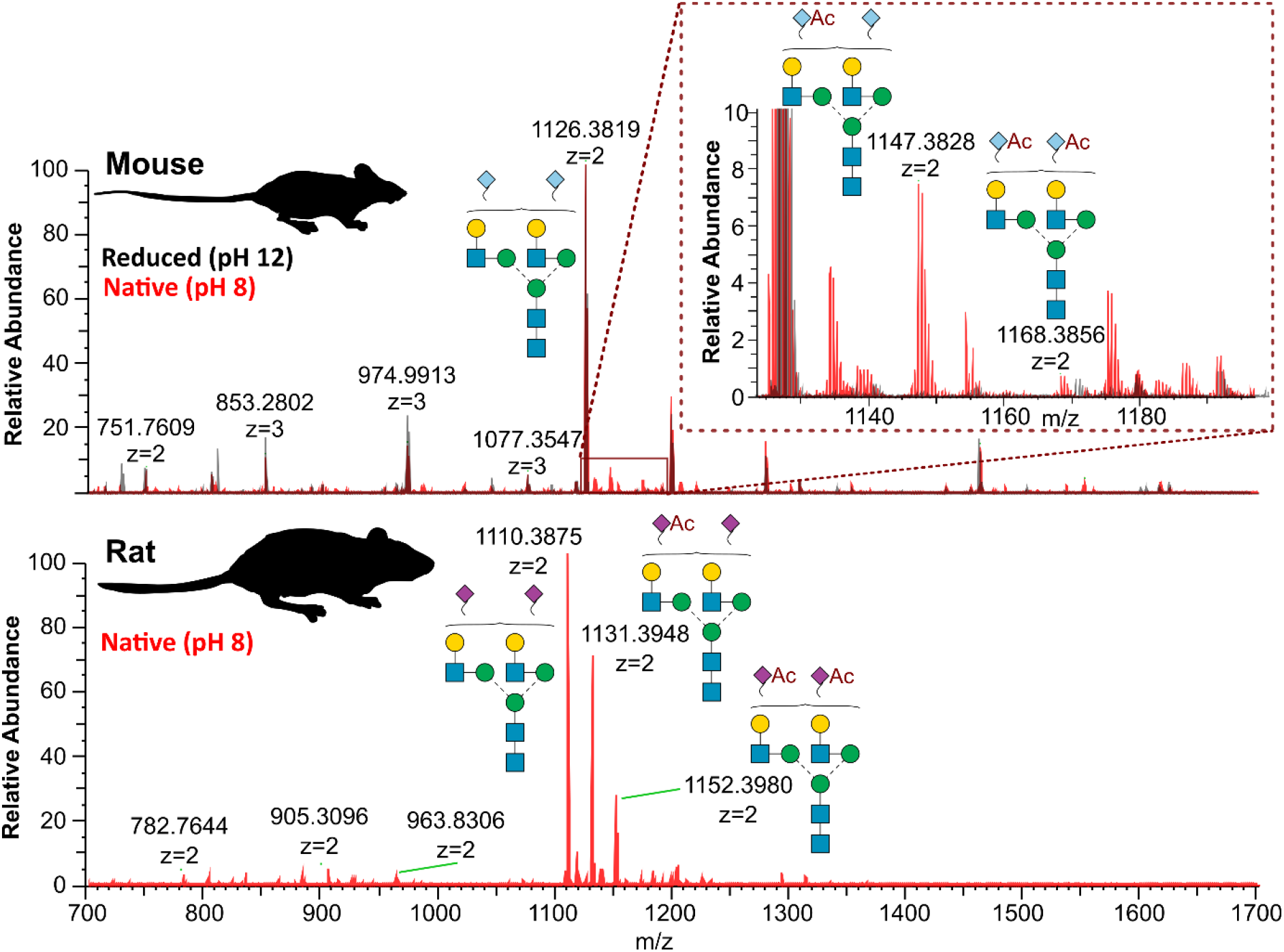
Native PGC-LC-MS workflows enable detection of *O*-acetylated *N*-glycans which are lost using glycan reduction. **Top** Mouse *N*-glycan mass profile for native and reduced conditions, inset 10-fold magnified spectrum revealing *O*-acetylation differences. m/z labels are for native *N*-glycans. **Bottom** Native conditions enable detection of *O*-acetylated NeuAc in rat serum.

Interestingly, similar *O*-acetylation had been observed by Helm *et al*.^20^, though at much lower levels, likely reflecting the high-pH reduction procedure used in that study. Although glycan release here was performed at pH 8, the extent of any pH-induced de-*O*-acetylation remains unknown. These findings contextualize previous glycan release strategies, including reductive or non-reducing β-elimination^21^, hydrazinolysis^22^, eliminative oximation^23^, and standard oxidative workflows^24^, which have not reported *O*-acetylated sialic acids due to their lability under alkaline conditions^10^.

Applying the same native workflow to rat serum uncovered pronounced *O*-acetylation of NeuAc. This suggests that this labile modification is common in mammalian sera and is best preserved under mild, non-reducing conditions. Furthermore, detection of *O*-acetylated NeuAc-containing *N*-glycans in rats, enables direct comparison to human sera, which is dominated by NeuAc. To make this method broadly applicable to sera derived from humans, LC-MS parameters for confident *O*-acetylated *N*-glycan detection were optimised.

To identify optimal fragmentation conditions for structural elucidation of *O*-acetylated sialylated glycans, resonant CID and beam-type CID were systematically compared using doubly acetylated NeuGc- and NeuAc-containing structures (**Figure 2A**). Both approaches generated high-mass fragments corresponding to the neutral loss of acetylated sialic acids; however, beam-type CID yielded more compositionally informative spectra. These included diagnostic ions at *m/z* 348 and 272 for acetylated NeuGc and NeuAc, respectively, and antenna fragments at *m/z* 713 and 697 retaining the *O*-acetyl group. Consequently, beam-type CID was selected for further optimisation.

**Figure 2.**
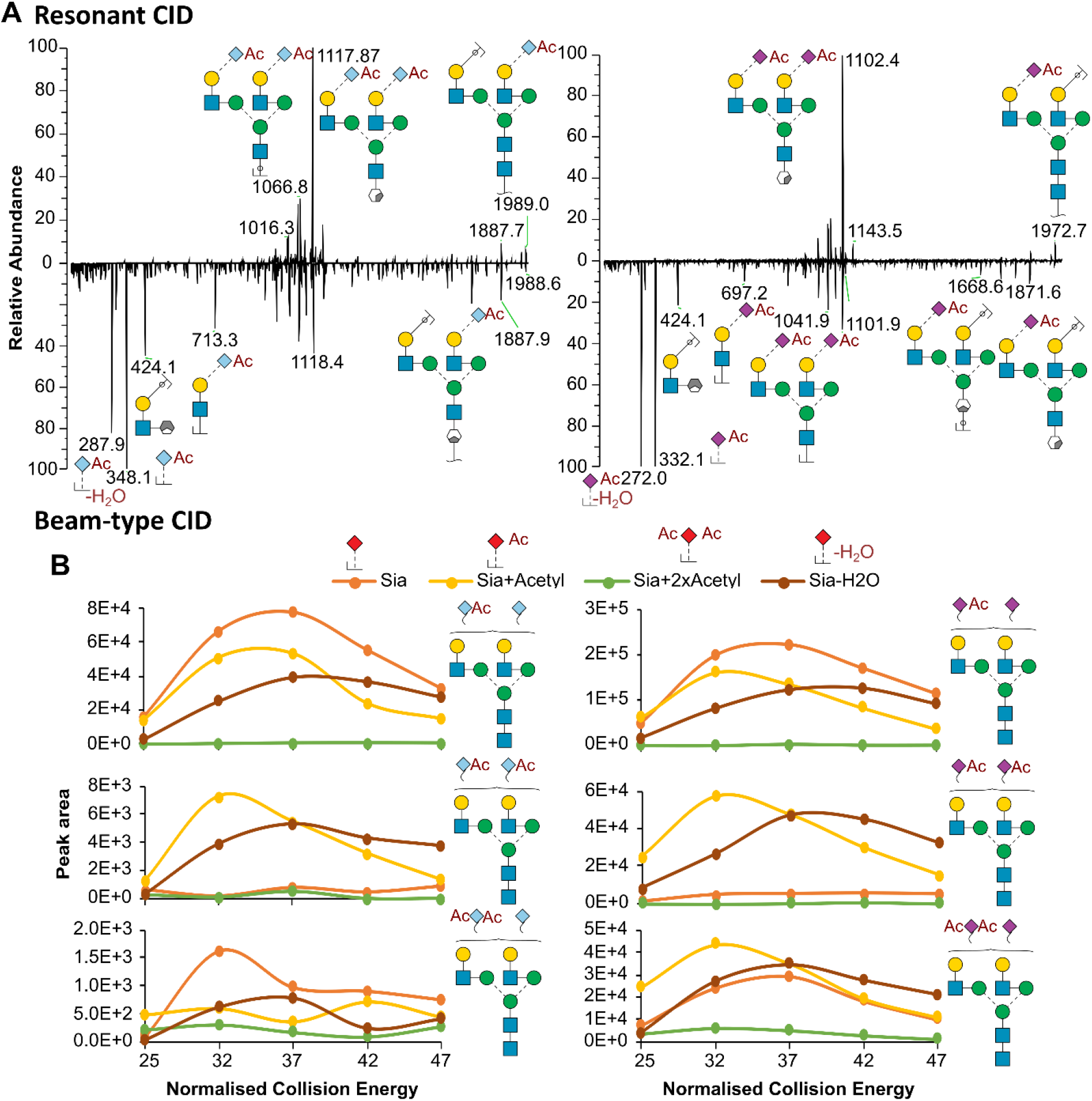
Fragmentation comparisons reveal optimal approaches and energy for *O*-acetylated *N*-glycan characterisation across NeuGc and NeuAc sialic acids. **A** Comparison between resonant CID and beam-type CID identifies beam-type CID as producing more informative product ions. **B** Collision energy optimisation identifies product ion ratios across different instrument energies

MS2 spectra were collected for the most abundant *O*-acetylated structures at multiple collision energies to establish a relationship between collision energy and product ion abundance. Increasing collision energy enhanced total product ion intensity, with a maximum observed at 37 NCE across all tested structures (**Figure 2B**). Notably, fragment ion ratios correlated with the degree of *O*-acetylation, with minimal unmodified sialic acid fragments detected in doubly *O*-acetylated glycans.

At higher energies, additional ions consistent with dehydration of sialic acid were observed. These diagnostic relationships between product ion intensity and *O*-acetylation state enable verification of proposed compositions and localisation of *O*-acetylation to individual monosaccharides.

Chromatographic behaviour on porous graphitised carbon (PGC) demonstrated that each additional *O*-acetyl group increased glycan hydrophobicity, resulting in later retention times relative to unmodified counterparts (**Figure 3A**). Beyond these expected shifts, *O*-acetylated species exhibited broadened and occasionally split peaks, suggesting multiple possible acetylation sites on the sialic acid residue and variable interactions between the *O*-acetyl group and the PGC stationary phase. Across all confidently assigned structures, including mono-, di-, tri-, and tetra-sialylated glycans, peak broadening was a consistent feature (**Figure 3B**). This behaviour implies that *O*-acetylation not only alters PGC retention but also introduces microheterogeneity within otherwise similar glycan species.

**Figure 3.**
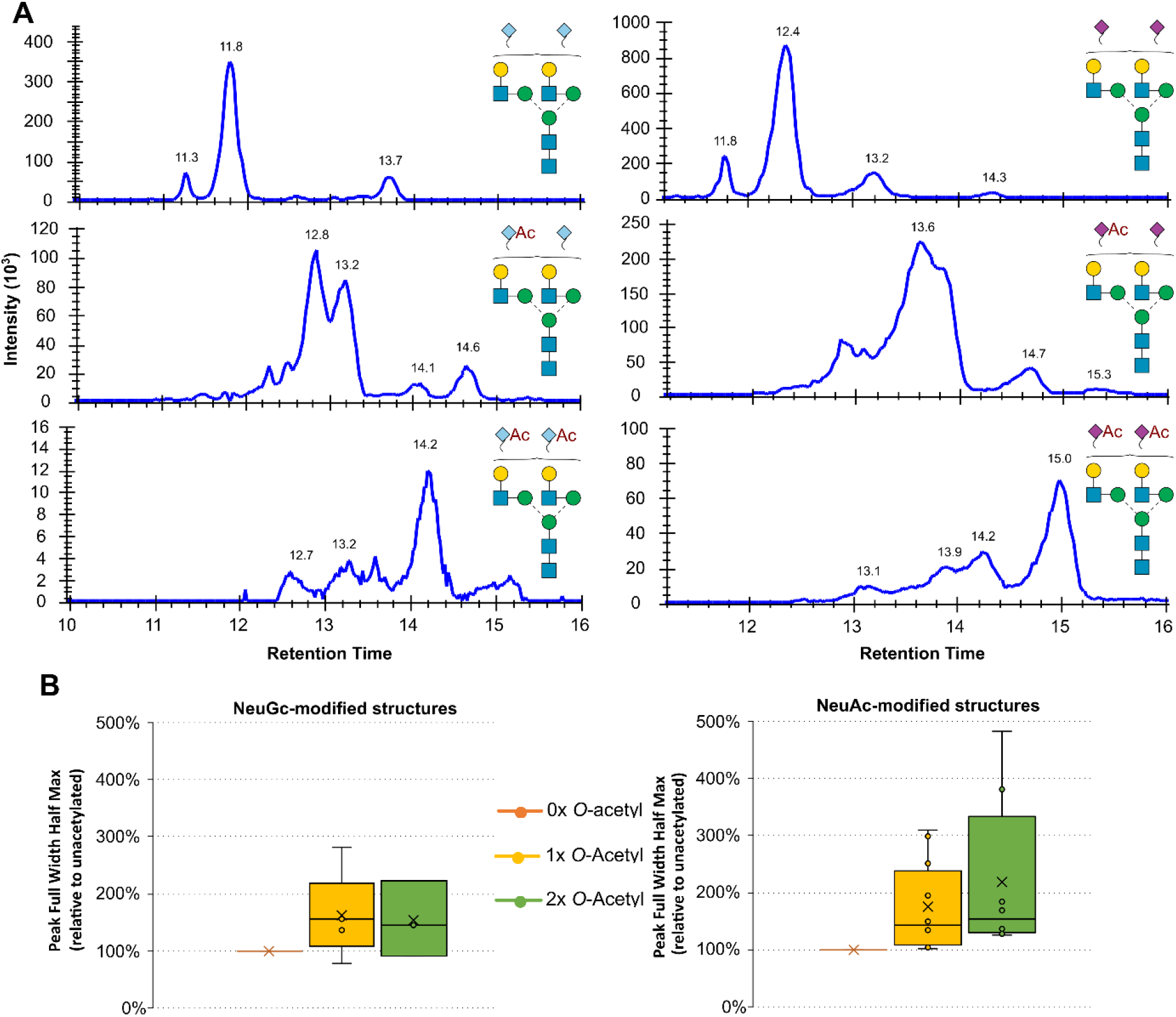
*O*-acetylation results in later eluting, broader peak shapes on PGC-LC. **A** Extracted ion chromatograms for unmodified, singly, and doubly *O*-acetylated *N*-glycans. **B** Box plot summarising peak width increases across all confidently identified *N*-glycan compositions

Vos *et al*.^25^ studied *N*-glycan *O*-acetylation orthogonally using ion mobility informed by synthetically generated fragments, enabling mapping of known structural epitopes to the *N*-glycans in contrast to this work. Similar to our study however, LC peak shapes on their ZIC-HILIC were observed to be unusually broad, indicating that this phenomenon is not stationary phase specific. These broads peaks could potentially be explained by *O*-acetyl migration, as observed and described by Ji *et al*.^26^, where minimal migration was only observed below pH 5, lower than that achievable at physiological pH. Regardless of the cause, these unusual chromatographic properties also provide an opportunity as a fingerprint of *O*-acetylation, as only *O*-acetylated structures were observed to have these broad peaks.

Leveraging these chromatographic and fragmentation insights, a checkpoint-based identification workflow was developed to improve confidence in *O*-acetylated *N*-glycan composition assignment (**Figure 4A**). Given that retention time mapping for *O*-acetylated glycans remains incomplete and MS^1^ data alone cannot resolve isomeric compositions, MS2 evidence was deemed essential for confident identification. Candidate structures were evaluated against theoretical isotopic distributions (idotp > 0.85), required to display characteristic diagnostic ions at expected ranked intensities, and verified to elute later than corresponding unmodified glycans. These metrics are summarised visually in **Figure 4B**. These quality-control criteria, when applied sequentially, substantially reduced the number of putative identifications, with only 5% of mouse and 3% of rat putative compositions meeting all thresholds (**Figure 4C**). While this stringent filtering reduces total identifications, it ensures robust, reproducible annotation of *O*-acetylated glycans.

**Figure 4.**
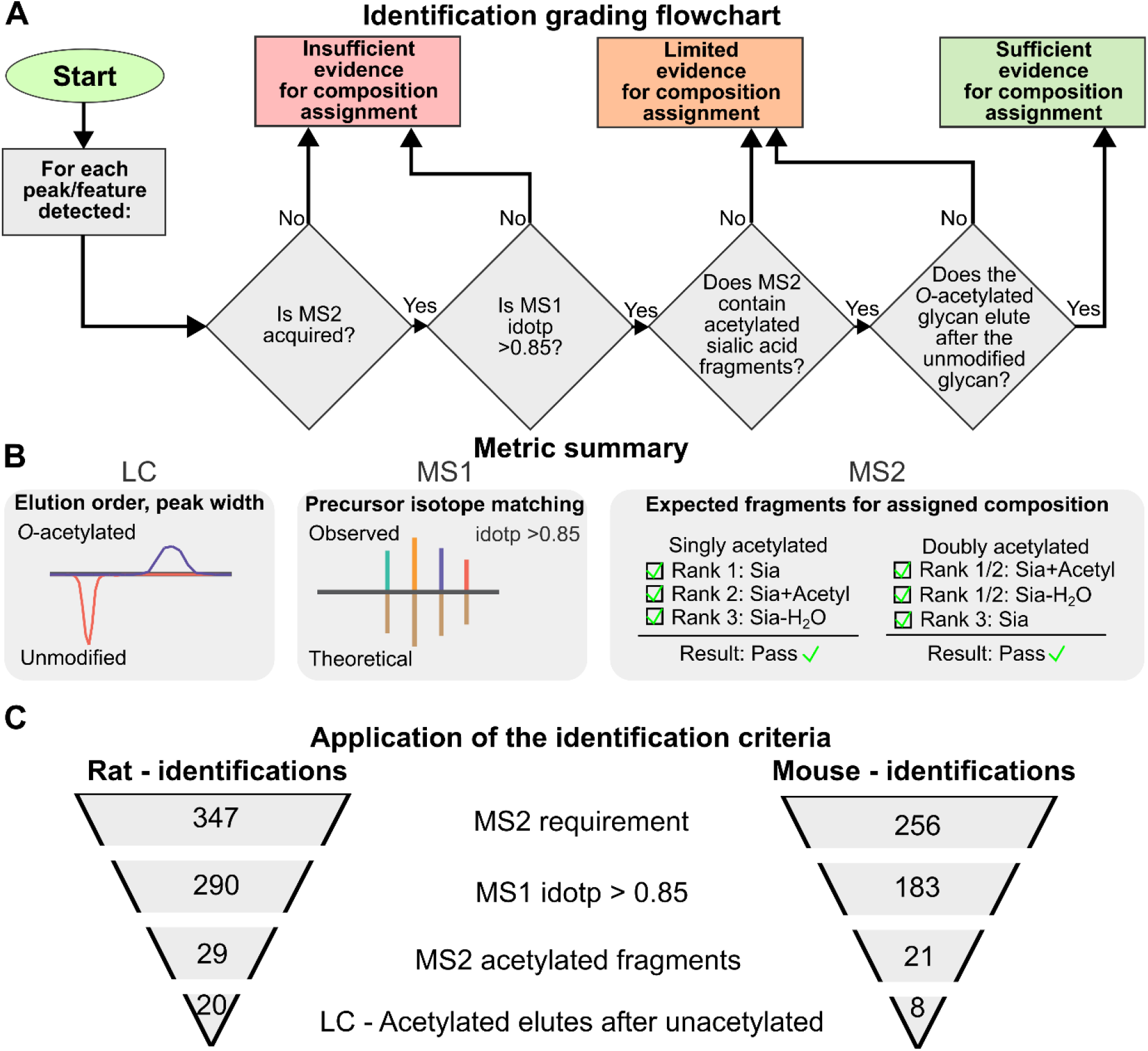
A checkpoint-based flow chart for *O*-acetylated *N*-glycan composition identification. **A** Graphical representation of the metrics used to verify identifications. **B** Flowchart for grading O-acetylated N-glycan identifications. **C** Impact of each criterion on number of composition identifications across rat and mouse sera *N*-glycans

Applying the optimised native PGC-LC-MS strategy to sera from mouse, rat, and human revealed striking interspecies differences in *O*-acetylation prevalence (**Figure 5**). Consistent with established reports^27–30^, mouse *N*-glycans were predominantly NeuGc-containing, whereas rat and human *N*-glycomes were primarily NeuAc-based. Quantitative analysis showed that 8.8% of mouse *N*-glycans contained at least one *O*-acetylated NeuGc residue, while rat sera exhibited extensive modification, with 53.4% of the total glycan peak area corresponding to *O*-acetylated NeuAc structures. Despite the detection of *O*-acetylated species for NeuGc and NeuAc in mouse and human sera, *O*-acetylation was undetected. Although *O*-acetylated sialic acids have been reported previously in monosaccharide-level analyses and individual glycoproteins such as erythropoietin^31^, the glycan species bearing such modifications in human serum remain unidentified. Together, these findings demonstrate that *O*-acetylation is a prevalent yet labile modification, variably expressed across rodents, and that native LC-MS workflows are essential for its reliable detection and structural characterisation.

**Figure 5.**
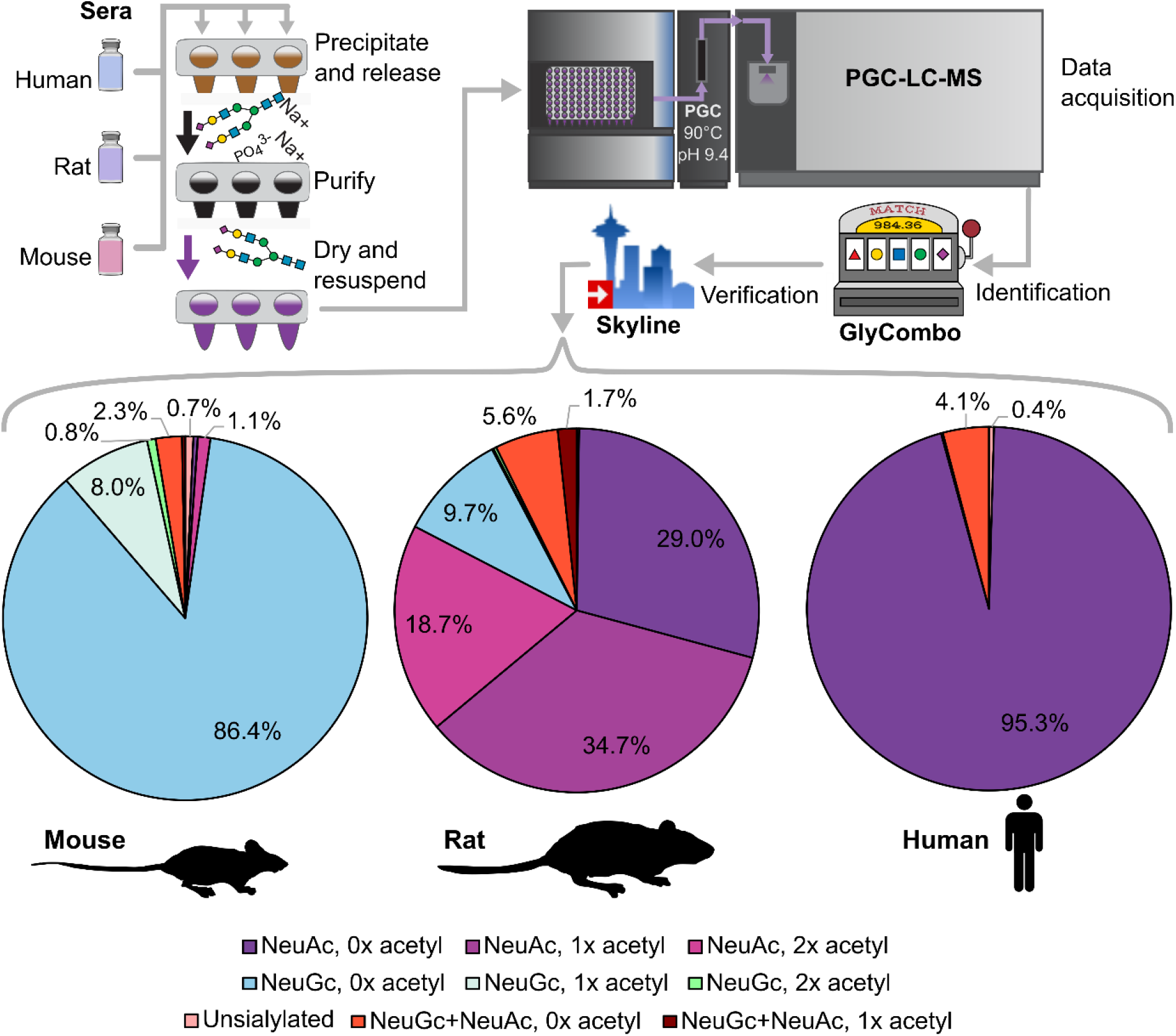
An optimised PGC-LC-MS acquisition strategy for native *N*-glycans reveals *O*-acetylation differences between NeuAc and NeuGc across mammalian model system sera, with an *O*-acetylated N-glycans undetected in human sera

Beyond glycomics, we reproduced the findings of Liu *et al*.^32^ which analysed complementary intact glycopeptides. Consistent with their observations, we detected greater than 50% sialic acid *O*-acetylation in rat serum and approximately 5% in mouse serum, with no evidence of *O*-acetylation in human serum. These results demonstrate strong concordance between released glycan and glycopeptide analyses, underscoring the reproducibility and cross-platform validity of native *O*-acetylation measurements. Our results also highlight the risks of optional high-pH steps in glycoproteomic workflows including TMT quenching with methylamine^33^ (pH 11) and high pH fractionation^34^ (pH 10).

## Conclusions

This study demonstrates that native PGC-LC-MS workflows enable the detection and characterisation of labile *O*-acetylated *N*-glycans that are lost under conventional high-pH derivatisation or reduction conditions. By maintaining the sample preparation pH at or below 8, we preserved *O*-acetylated NeuGc and NeuAc species in mouse and rat sera, revealing that this modification is more prevalent than previously recognised. Comparative fragmentation analysis established beam-type collision-induced dissociation as the optimal approach for structural elucidation, providing diagnostic product ions that verify both the presence and degree of sialic acid *O*-acetylation. Collision energy optimisation further refined these measurements, identifying 37 normalized collision energy as the most informative setting for *O*-acetylated sialylated glycans.

Chromatographic evaluation showed that each additional *O*-acetyl group increased glycan hydrophobicity, producing later elution and broader peak shapes indicative of multiple acetylation sites and conformational heterogeneity. Integrating these chromatographic and spectral characteristics, a checkpoint-based identification workflow was established, combining isotopic accuracy, diagnostic fragment detection, and retention-time shifts to ensure confident assignment of *O*-acetylated structures. Application of this framework reduced false identifications while maintaining robust reproducibility across species.

Quantitative profiling revealed distinct species-specific *O*-acetylation patterns: extensive modification in rat serum (53.4%), moderate levels in mouse serum (8.8%), and none detectable in human serum. Together, these findings confirm that *O*-acetylation is a widespread but labile modification that varies across mammalian systems. The optimised native PGC-LC-MS strategy provides a reliable foundation for future studies exploring the structural diversity, biological significance, and regulatory mechanisms of glycan O-acetylation.

## Author contributions (with CRediT details)

CRediT: Conceptualization: CA; Data curation: CA; Formal Analysis: CA; Funding acquisition: CA; Investigation: CA; Methodology: CA; Project administration: CA; Visualization: CA; Writing – original draft: CA; Writing – review & editing: CA

## Data availability

Raw files, and Skyline assays are available on Panorama Public^35^ at: https://panoramaweb.org/NGOacetyl.url and GlycoPost^36^ at: https://glycopost.glycosmos.org/entry/GPST000636. GlyCombo output, collision energy optimisation results, and *N*-glycan quantitation values are available in a Supplementary excel spreadsheet.

## Acknowledgements

This research was facilitated by access to Sydney Mass Spectrometry, a core research facility at the University of Sydney.

